# Inhibition of cAMP signaling prevents congenital heart defects counteracting oxidative stress in Pde2A deficient embryos

**DOI:** 10.1101/2023.05.29.542779

**Authors:** Silvia Cardarelli, Martina Biglietto, Tiziana Orsini, Valentina Fustaino, Lucia Monaco, Ana Gabriela de Oliveira doRego, Francesca Liccardo, Silvia Masciarelli, Francesco Fazi, Fabio Naro, Luciana De Angelis, Manuela Pellegrini

## Abstract

**Background:** Phosphodiesterases (PDEs) are the enzymes that hydrolyze cyclic nucleotides (cAMP and cGMP) playing a key role in the homeostasis of these two second messengers. PDE2A is a dual-specific PDE that breaks down both cAMP and cGMP and can be activated by cGMP. It appears peculiar that the Pde2A-deficient (*Pde2A^-/-^*) mouse model is embryonically lethal, likely due to a strongly reduced size of liver and to a severe anemia. In addition, the heart of *Pde2A^-/-^* embryos shows ventricular and atrial septum defects, hypertrabeculation, heart dilatation and non-compaction defects. We recently highlighted a direct relationship between Pde2A impairment, consequent increase of cAMP and the onset of mouse congenital heart defects (CHDs), however the molecular mechanisms underlining the heart defects remain unknown.

**Methods:** Transcriptome analysis of *Pde2A^-/-^* embryonic heart was performed by RNA sequencing and the most altered genes were also analyzed by quantitative real time PCR. In vivo treatment with drugs acting on cAMP signaling (Metoprolol and H89) and oxidative stress (N-Acetyl-Cysteine, NAC) were carried out on pregnant *Pde2A^+/-^* female. Histological, biochemical, and molecular analyses were then performed on embryonic hearts.

**Results:** We found a significant modulation of more than 500 genes affecting biological processes involved in the immune system, cardiomyocyte development and contractility, angiogenesis, control of gene transcription and oxidative stress in hearts from *Pde2A^-/-^* embryos. Metoprolol and H89 administration were able to prevent heart dilatation and hypertabeculation in *Pde2A^-/-^*embryos. Metoprolol was also able to partially impede heart septum defect and oxidative stress at tissue and molecular levels. Partial rescue of cardiac defects was observed by using the antioxidant NAC, indicating oxidative stress like one of the molecular mechanisms underpinning the CHDs.

**Conclusions:** We identified specific biological processes, molecules and cell signaling that can be targeted by selected drugs with consequent beneficial effects for cAMP-dependent CHDs.

**Novelty and Significance:** *What is Known?:*

- Congenital Heart Defects are the most frequent heart birth defects including septal defects, hypertrabeculation and non-compacted myocardium.
- Pde2A hydrolyses the cAMP and cGMP second messengers.
- *Pde2A*-deficient mice are embryonic lethal and show cAMP-dependent Congenital Heart Defects.

*What New Information Does This Article Contribute?:*

- We identified several novel pathways altered in hearts of *Pde2A^-/-^* embryos.
- We demonstrated that drugs lowering cAMP levels rescued specific CHDs in *Pde2A^-/-^* embryos.
- We discovered that antioxidants are beneficial for CHDs in *Pde2A^-/-^* embryos. The significance of this work relay in molecular discoveries and pharmacological approaches to treat CHDs by using a mouse model that recapitulate the major congenital heart defects. Among the pathways involved in specific defects associated with CHDs, the transcriptome analysis revealed an impairment of genes of the immune system, cardiomyocyte development and contractility, angiogenesis, control of gene transcription and oxidative stress in *Pde2A^-/-^* hearts. The scientific community will have open access to the RNA-seq data that can be utilized to further understand the congenital cardiac pathology and clarify the molecular implication in selected defects such as septal and ventricular wall defects. Up to date CHDs, when possible and if identified in time, are mostly treated trough surgery. The identification of drugs blunting the cAMP signaling response or reducing oxidative stress pathways will be useful for setting therapeutic approaches to alleviate CHDs.

## INTRODUCTION

Among phosphodiesterases (PDEs), the specific enzymes that hydrolyze cyclic nucleotides (cAMP and cGMP), PDE2A is unique, because the knockout mouse model is embryonically lethal probably due to an extreme reduced liver size, impairment of the hepatic niche and elevated anemia^1, 2^.

We recently reported that hearts of *Pde2A^-/-^* embryos and hearts of embryos treated with a Pde2A inhibitor show ventricular and atrial septum defects, hypertrabeculation, heart dilation and non-compaction defects^3, 4^.

We also demonstrated a direct relationship between Pde2A deficiency, the consequent increase of cAMP and the onset of mouse congenital heart defects (CHDs). Indeed, during cardiogenesis, cAMP has been shown to significantly transform fetal cardiomyocytes into cardiac pacemaker-like cells increasing gap-junction markers^5, 6^ to promote phosphorylation of targets such as L-type calcium channel via Protein Kinase A (PKA) and to decrease its concentration during heart development^7, 8^. Pde2A is localized at the plasma membrane, in the cytoplasm and in the nucleus of neonatal murine cardiomyocytes.^4, 9^

PDE2A2 isoform was also described to be localized to the mitochondrial matrix and its inhibition to stimulate oxidative phosphorylation^10^. More recently, cAMP/PKA signaling domains localized at mitochondrial membranes and regulated by PDE2A2 were described to be involved in mitochondrial morphology change and apoptosis^11^. Because of this specific localization and function, we might hypothesize a contribution of PDE2A in reactive oxygen species (ROS) production. Indeed, ROS have traditionally been regarded as by-products of aerobic metabolism, and mitochondrial respiration is the major intracellular source of accidental ROS production^12^.

The stimulation of β-1 adrenergic receptor (β1-AdR) is the primary source of cAMP in the heart trough adenylyl cyclase (AC) activation^13^. Several uncontrolled and controlled clinical studies provided the first evidence that Metoprolol, a selective β1-adrenoceptor blocking agent (β-blocker), had beneficial clinical and haemodynamic effects in patients with heart failure due to idiopathic dilated cardiomyopathy (DCM) or ischaemic heart disease or ischaemic heart disease^14–18^. Metoprolol was able to extend survival in DCM mice and to prevent cardiac remodelling and dysfunction^19–20^.

The cAMP target PKA plays several functions in the normal heart such as contraction, metabolism, and gene transcription regulation. However, increase of PKA activity and protein level, were described to aggravate heart failure^21–23^. PKA activators induced ROS and consequent electrical disturbance in heart failure, whereas PKA inhibition with H89 prevented it in humans^23^. H89 administration resulted also in attenuation of dilated cardiomyopathies and prevention of heart failure^24^. Moreover, it has been demonstrated to decrease cell death in heart disease and to improve cardiac function after myocardial infarction in mice^25^.

Here, investigating the transcriptome of Pde2A deficient embryonic heart by RNA sequencing (RNA-seq) analysis, we found a significant modulation of more than 500 genes associated to biological processes such as immune response, cardiomyocyte development and contractility, angiogenesis, control of gene transcription and oxidative stress. Testing pharmacological treatments that blunt cAMP/PKA signaling we observed that Metoprolol and H89 can robustly prevent CHDs in Pde2A deficient embryos such as N-Acetyl-Cysteine (NAC) antioxidant administration. Moreover, among the pathways severely up-regulated, we found Metoprolol treatment able to re-establish the expression levels of oxidative-stress-related genes and NAC to reduce in vivo heart oxidative stress.

## METHODS

### Mouse breeding and in vivo treatments

*Pde2A^+/−^* mice (B6; 129P2-Pde2A < tm1Dgen>/H; EM: 02366) were obtained from EMMA (UK). Timed mating was set up between *Pde2A^+/−^* females and *Pde2A^+/−^* males. Genotyping for Pde2A was performed as previously reported.^3^ To test if pharmacological treatments, that affect directly or indirectly cAMP, can rescue cardiac malformations of *Pde2A^-/-^* mice we chose two drugs that inhibit β1-adrenergic receptor signaling (Metoprolol) and PKA activity (H89), respectively. The drugs were administered every day starting at E5.5 from the plug until E13.5. Embryos were collected at E14.5. Metoprolol (Sigma-Aldrich) was administered by gavage at 10 mg/kg dissolved in drinking water^19, 26^. H-89 (Sigma-Aldrich) was administered intraperitoneal at 1 mg/kg^27, 28^.

NAC (Sigma-Aldrich) was administered at 1 g/kg/day to *Pde2A^+/-^*females in drinking water starting the first day of the plug and until day E14.5^29^. Hearts of the resulting E14.5 embryos were isolated in Hanks balanced solution as previously described^3, 30^ and processed for following analyses.

All our experimental animal procedures were conformed to the Directive 2010/63/EU and conducted with the approval of the Italian Ministry of Health 919/2020-PR in date 28/9/2020.

### Micro-CT Imaging and Volume Measurements

Imaging specimen preparation: embryos were fixed in 10% neutral buffered formalin (Bio-Optica) or 4% paraformaldehyde (PAF, Sigma-Aldrich) overnight at room temperature (RT) and embedded in paraffin following standard procedures.

Micro-CT scanning: Computed tomography images were acquired by a high-resolution 3D micro-CT imaging system (Skyscan 1172G Bruker, Kontich—Belgium), using a L7901-20 Microfocus X-ray Source (Hamamatsu). The acquisition of volumes was performed in 1.5 mL micro-tubes, with a camera pixel/size of 7.9 µm, camera binning 2×2, tube voltage peak of 39 kV, tube current of 240 µA, exposure time of 450 ms. Reconstructions of tomographic datasets were performed using built-in NRecon Skyscan Software (Version:1.6.6.0; Bruker). The 3D volumes were analyzed using 3D Visualization Software CTvox v. 2.5 (Bruker). Tissues segmentation: Manual image-by-image segmentation aimed at calculation of heart volume was applied, using Bruker micro-CT Analyzer Version 1.13 software. A histological atlas of mouse development^31^ was used for guidance to accurately identify, demarcate, and segment each embryo (N = at least 3) in a specific VOI (Volume of Interest) for automated volume measurements^31, 32^.

### Hematoxylin and Eosin staining and Immunofluorescence

Embryos at E14.5 were fixed in 10% neutral buffered formalin (Sigma-Aldrich) overnight at RT and included in paraffin (Sigma-Aldrich) following standard procedures. Serial paraffin sections of whole embryos (5 μm of thickness) were obtained, deparaffinized and stained with hematoxylin and eosin (Sigma-Aldrich). Wall and trabeculae thickness, contralateral axis length, trabeculae number were evaluated as previously described^3^. All histological analyses were performed in a blinded way by two different authors.

For immunofluorescence, 5μm E14.5 paraffine sections underwent deparaffinization and microwave antigen retrieval in pH 6.0 sodium citrate with 0.05% Tween-20 solution (Sigma-Aldrich), followed by gradual chilling. Samples were permeabilized with 0.2% NP40 (Sigma-Aldrich) in PBS for 30 minutes and blocked with 1% bovine serum albumin (Sigma-Aldrich) in Phosphate Buffered Saline (PBS) for 1 hour at room temperature. Sections were incubated overnight at 4°C with anti-Endomucin antibody (Ab106100, Abcam). After 3×5min washes with PBS, sections were stained for 1 hour at room temperature with anti-rat secondary antibody (Alexa Fluor 488; Thermo Fisher Scientific) followed by 2 minutes of Sudan Black (Sigma-Aldrich; 0,03gr in 10ml di ET-OH70%) to reduce the background and nuclei were counterstained with Hoechst (Sigma-Aldrich) for 5 minutes. Slides were mounted with Fluoromount Aqueous Mounting Medium (Sigma-Aldrich).

### RNA-sequencing and qRT-PCR

Total RNA was isolated from embryonic cardiac tissue using Total RNA Purification kit (Norgen) according to manufacturer’s instruction. For RNA-sequencing, the integrity and purity of the RNA was initially checked at the Bioanalyzer (Agilent 2100 bioanalyzer) by using RNA 6000 Nano Agilent kit. and the RNA seq analysis was done at EMBL service facility, Heidelberg. mRNA-Seq libraries were prepared from total RNA samples treated with DNase and single-end sequenced on the specified Illumina sequencing instrument. RNA-seq data are available online at geo platform.

For quantitative RT-PCR, mRNA was reverse transcribed to cDNA through Maxima H minus reverse transcriptase (Thermo Fischer Scientific). qRT-PCR reaction was carried out by using PowerUp SYBR green Master Mix (Thermo Fisher Scientific). Target transcripts were analyzed using Applied 7500 sequence detector system (Applied Biosystems, Carlsbad, CA, USA).

For the quantification analysis, the comparative threshold cycle (Ct) method was used. The Ct values of each gene were normalized to the Ct value of β-Act in the same RNA sample. The gene expression levels were evaluated by fold change using the equation 2^-ddCt^. Primers used in mRNA analyses are reported in Table S1.ù

### cAMP and cGMP assay

Embryonic hearts were homogenized and sonicated in 10 volumes of 0.1 M HCl with 0.1% Triton X-100 and processed with the Direct cAMP/cGMP ELISA kit (Enzo) following the manufacture’s instruction. Heart samples and standards were acetylated before being loaded into the plate by using Acetic Anhydride and Triethylamine 1:2. Samples were incubated over night at 4°C on a plate shaker. A substrate solution was added and incubated for 1 hour at room temperature. The reaction was stopped, and samples were read at the spectrophotometer at an optical density of 405nm. Results are referred to a cAMP and cGMP Standard Curves performed together with the cAMP and cGMP Assay.

### Reactive Oxidative Stress detection

Oxidative stress was quantified by measuring total reactive oxygen species (ROS) using 2μM 2’,7’-dichlorodihydrofluorescein diacetate (CM-H2DCFDA; Sigma-Aldrich) staining and FACS analysis. Briefly, heart embryos were dissected, washed in PBS and dissociated with 5 mg/ml Collagenase Type II (Sigma-Aldrich) at 37 °C for 30 min. After washing with PBS, cells were resuspended in PBS and stained with CM-H2DCFDA for 30 min at 37°C in the dark. At the end of the incubation, to remove dye, cells were centrifuged and washed with PBS. The final cell suspensions were filtered through a 70 μm strainer, stained with Sytox Blue Dead Cell Stain (Thermo Fisher Scientific) and analysed on a Cytoflex (Beckman Coulter). Flow cytometry data were analysed with CytExpert software.

### Statistical analyses

All data are expressed as mean ± SEM and analyzed with Student t-test with two tails or ANOVA-two way with Tukey correction. Differences were considered significant if * P<0.05.

## RESULTS

### RNA-seq analysis reveals alteration of pathways involved in immune system, angiogenesis, oxidative stress, gene transcription and cardiogenesis

We previously showed downregulation of critical genes involved on cardiac development in *Pde2A* knockout embryos compared to wild types^3^. To deeper delineate the global transcriptional profile of the two genotypes, RNA-seq was performed on isolated hearts from *Pde2A^+/+^* and *Pde2A^-/-^*embryos at E14.5.

Principal component analysis (PCA) indicated that *Pde2A^-/-^* was consistently different from *Pde2A^+/+^*, with the first component showing more than 50% of the global variance (Figure 1A). The extensive alteration in the expression profile between *Pde2A^-/-^* and *Pde2A^+/+^*was also displayed by the heatmap (Figure 1B): a stringent analysis (fold change ≥2; p value < 0.05) found 55 downregulated genes and 460 upregulated genes in the *Pde2A^-/-^* with respect to *Pde2A^+/+^*embryonic hearts (Table S2). A subsequent gene ontology (GO) analysis evaluated by the DAVID functional enrichment on biological processes suggested that upregulated genes were involved in pathways such as immune system and inflammatory response, chemokine-mediated signaling, angiogenesis, cellular response to hypoxia and positive regulation of ERK1 and ERK2 cascades (Figure 1C). The differentially expressed genes were investigated using the Gene Set Enrichment Analysis (GSEA) and compared to gene sets from Molecular Signatures Database (MSigDB) to characterized different cellular populations enriched in *Pde2A^-/-^* hearts. In line with our previous data, the cardiomyocyte lineage negatively correlated with *Pde2A^-/-^*(Figure 1D), whereas the other lineages such as endothelial, fibroblast, smooth muscle cells, macrophages, are strongly positively correlated with *Pde2A^-/-^*(Figure S1).

**Figure 1.**
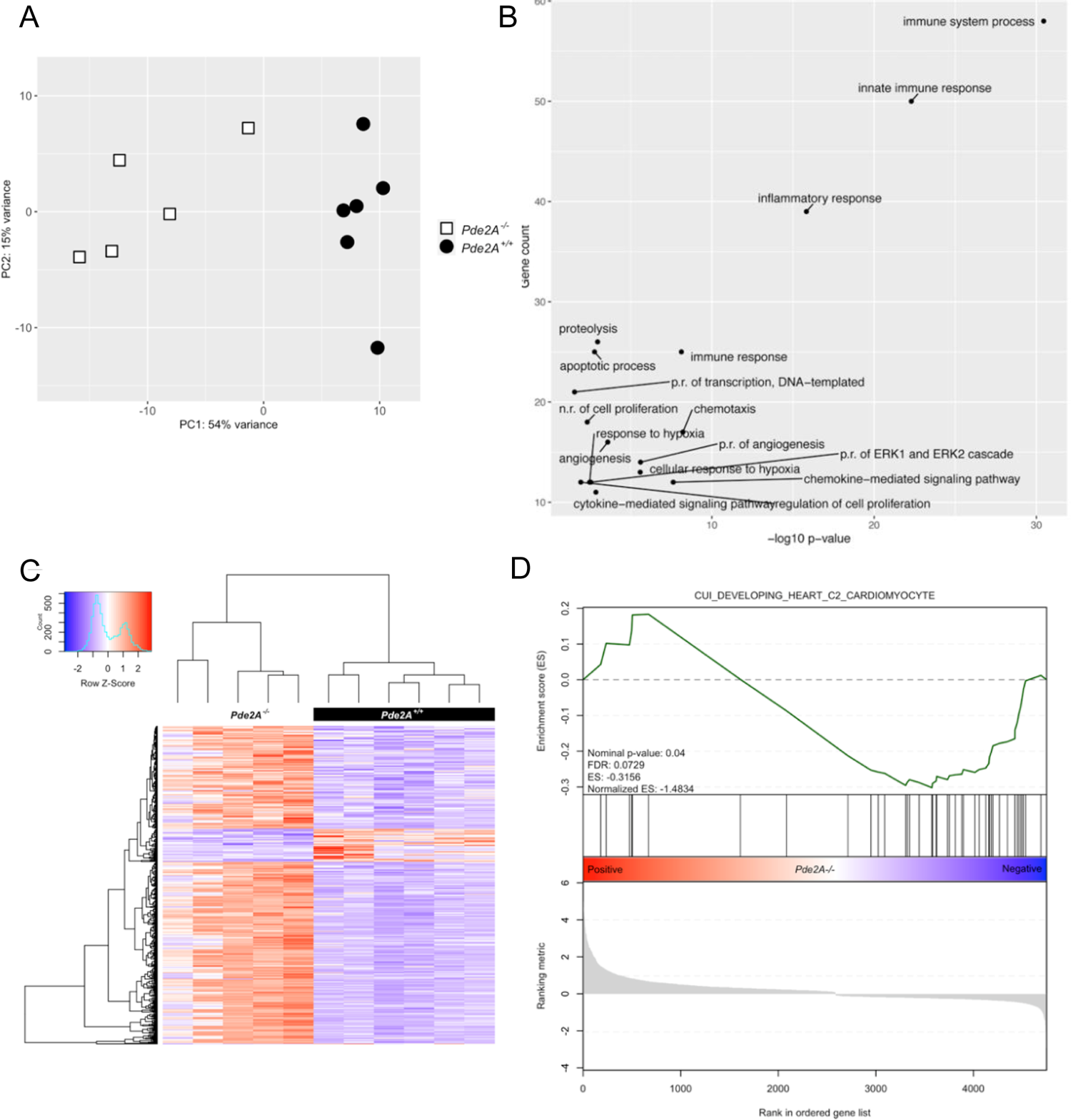
RNA-seq analysis reveals gene expression dysregulation in hearts of E14.5 *Pde2A^-/-^*. A) Plots of samples on first (PC1) and second (PC2) principal components of RNA-seq data. The PC percentage of variance is reported. B) Heatmap representation of 515 differentially expressed genes considering a log2-fold change >+1 and <-1. p adj < 0.05. C) GO-Enrichment in Biological Processes (BP) of genes significantly deregulated - p value<0.05 and gene count > 10. D) Gene Set Enrichment Analysis (GSEA) compared to gene sets from Molecular Signatures Database (MSigDB); n=6 *Pde2A^+/+^* and n=5 *Pde2A^-/-^* hearts.

Some genes, associated to enriched pathways found in DAVID analysis, were validated by qRT-PCR (Figure 2). We selected genes linked to the immune system/inflammation (Figure 2A), genes involved in angiogenesis and hypoxia (Figures 2B-C), genes involved in survival, proliferation, and transcription (Figures 2D-E). Some genes, such as *Ccl2, Vegfa*, *Il1a, Fgf23,* were robustly upregulated and expressed in different biological processes. Real-time PCR fully confirmed the RNA-seq analysis. A deeper investigation of RNA-seq results led to the hypothesis that also several genes involved in oxidative stress were altered in *Pde2A^-/-^* embryonic hearts (Table S2). Real-time PCR analysis revealed that most of them were upregulated in knockout hearts at E14.5 (Figure 2F). These results indicated that in *Pde2A^-/-^* hearts there is a high level of inflammation, oxidative stress, angiogenesis, and a reduction of cardiomyocyte differentiation, at least at transcriptional level.

**Figure 2.**
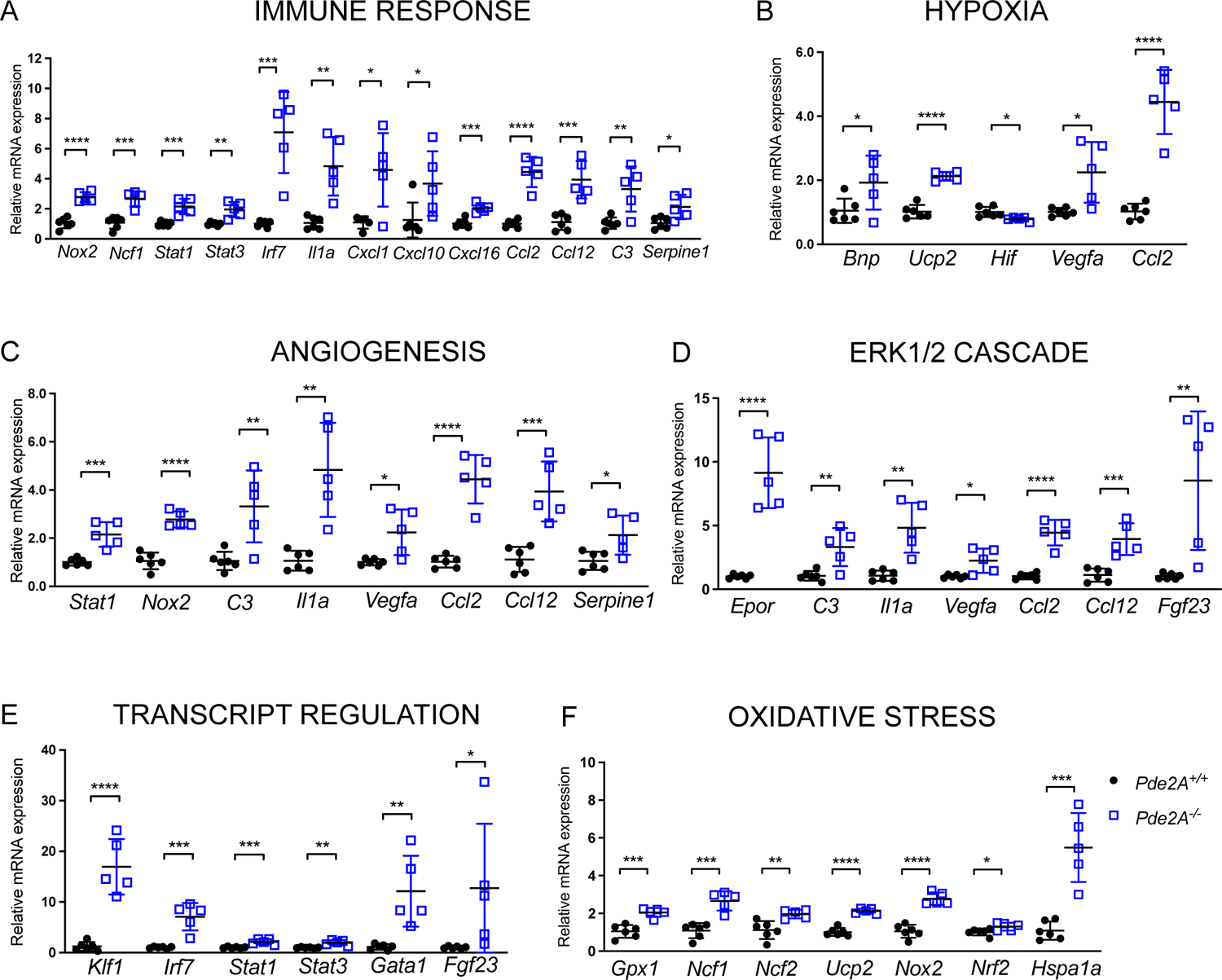
Real Time PCR confirms up-regulation of selected genes from biological processes. Histograms of relative mRNA expression of genes up-regulated in RNA-seq analysis and enriched in biological processes. Each dot represents an embryo; n=5 *Pde2A^+/+^* and n=5 *Pde2A^-/-^* hearts; unpaired Student’s t-Test * P≤0.05, ** P≤0.01, *** P≤0.001, **** P≤0.0001.

### Metoprolol treatment prevents CHDs of *Pde2A^-/-^* embryos

PDE2A can hydrolyze both cAMP and cGMP and, moreover, its catalytic activity increases in response to cGMP, which allosterically binds to the PDE2A GAF-B domain.

We measured the cyclic nucleotide levels in embryonic hearts from *Pde2A^+/+^* and *Pde2A^-/-^* and we observed that cAMP level is almost doubled in knockout compared to wild-type hearts at E14.5 (Figure 3A) as previously published^3^ whereas negligible differences were found in cGMP content (Figure 3B). Starting from these data, we decided to focus on pharmacological treatments that directly impact on cAMP signaling to rescue cardiac malformations in *Pde2A^-/-^*embryos. Metoprolol, a specific antagonist of β1-adrenergic receptor and consequently of cAMP synthesis, was administered after implantation to pregnant female mice every day, starting at E5.5 from the plug until E13.5, the day before sacrifice (Figure 3C). The level of cAMP was similar in *Pde2A^+/+^* and *Pde2A^-/-^* hearts after Metoprolol treatment validating the treatment efficacy (Figure 3D).

**Figure 3.**
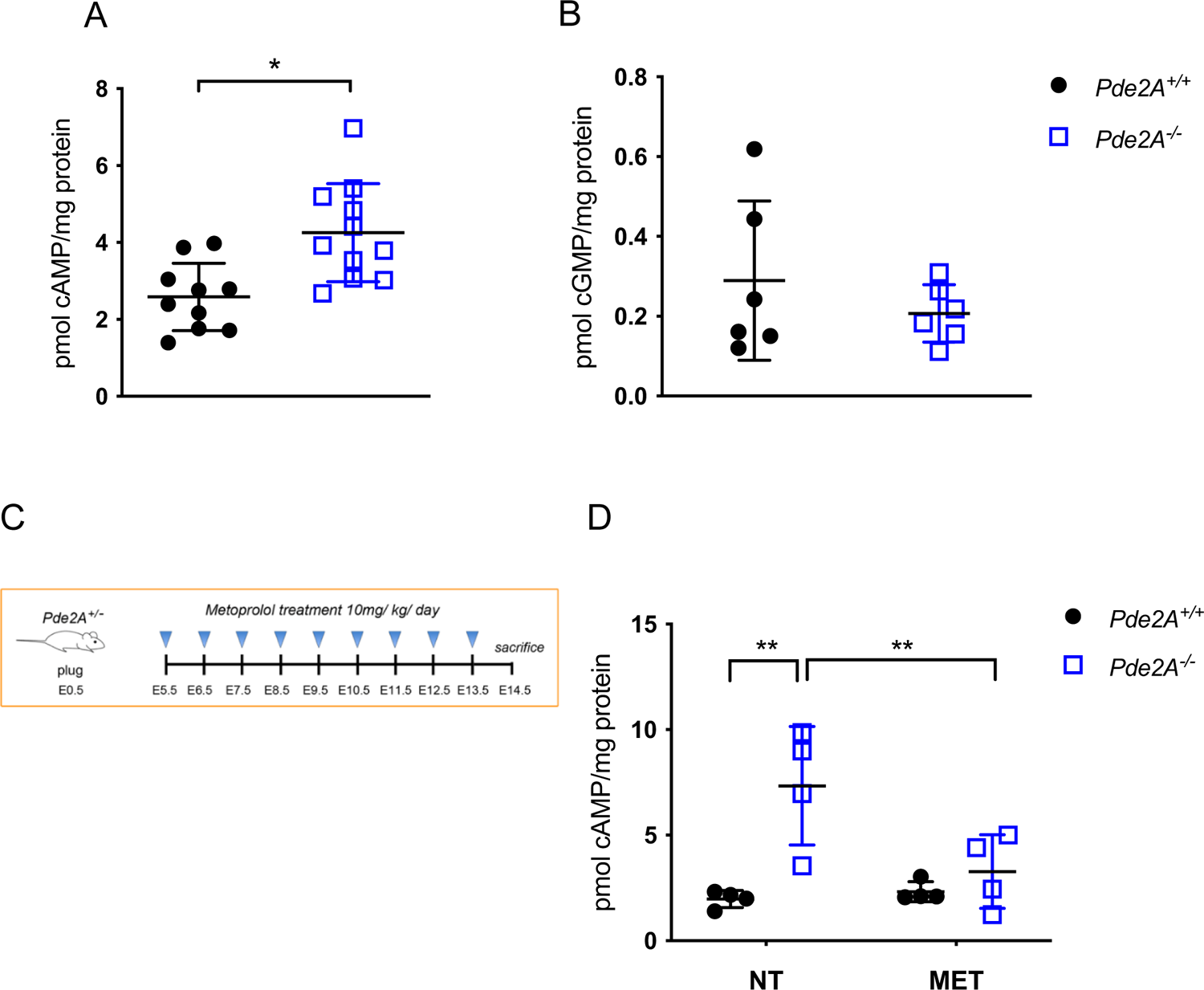
cAMP but not cGMP levels are modified in hearts from *Pde2A^-/-^* embryos. A, B) cAMP and cGMP levels in *Pde2A^+/+^* and *Pde2A^-/-^* hearts. At least n=6 *Pde2A^+/+^* and n=7 *Pde2A^-/-^* hearts; unpaired Student’s t-Test, * P≤0.05. C) Scheme of Metoprolol treatment. D) cAMP level in *Pde2A^+/+^* and *Pde2A^-/-^* in hearts of Metoprolol treated (MET) or not treated (NT) embryos. n=4 for each condition and genotype; ANOVA two-way was used to compare *Pde2A^-/-^* versus the relative *Pde2A^+/+^* and *Pde2A^-/-^* versus *Pde2A^-/-^* MET samples. ** P≤0.01, not significant values were not shown.

Morphometric analysis showed that Metoprolol did not affect hearts of *Pde2A^+/+^* embryos (Figure 4 and Figure 5). Moreover, morphological examination of Metoprolol-treated E14.5 *Pde2A^-/-^* embryos showed, like *Pde2A^-/-^* untreated embryos^3^ (Figure S2A), anemia, hemorrhages, and reduced liver size (Figure 4A and Figure S2A). However, micro-CT analysis revealed similar parameters between *Pde2A^-/-^* embryonic hearts treated with metoprolol and *Pde2A^+/+^* hearts (Figures 4B-E). Total heart volume, atrial volume, and ventricular volume normalized to embryos volume did not show difference in wild-type versus knockout treated embryos contrary to wild-type versus knockout untreated embryos (Figure 4C-E). Moreover, from micro-CT examination half of the *Pde2A^-/-^* treated embryos had no interventricular septum defects (Fig. 4B and Movie S1-S3). Morphological and morphometric analyses of the hearts were then performed on hematoxylin and eosin-stained sections of paraffin embedded E14.5 embryos. Contrary to untreated embryos, no differences in the contralateral axis between *Pde2A^+/+^*and *Pde2A^-/-^* treated hearts were observed (Figures 5A-B), confirming the ability of Metoprolol to prevent cardiac enlargement. Similarly, the hypertrabeculation of *Pde2A^-/-^* hearts was improved by Metoprolol administration and wild-type and knockout embryos showed similar thickness and number of cardiac trabeculae, respectively (Figures 5A-C-D). Furthermore, immunofluorescence for endomucin, that marks the endocardium, revealed trabecular network reconstitution after Metoprolol treatment in *Pde2A*^-/-^ embryos (Figure S3). However, a significant thinning of ventricular myocardium wall persisted in *Pde2A^-/-^*heart after Metoprolol treatment, even though a trend of thickness increase was observed (Figures 5A-E). Hematoxylin and Eosin staining confirmed that half of treated *Pde2A*^-/-^ hearts did not show ventricular septum defects (Figure 5A). These results suggest that Metoprolol, reestablishing the cAMP levels, can partially revert *Pde2A^-/-^* cardiac defects, probably acting on some of the pathways specifically controlled by the Pde2A/cAMP system.

**Figure 4.**
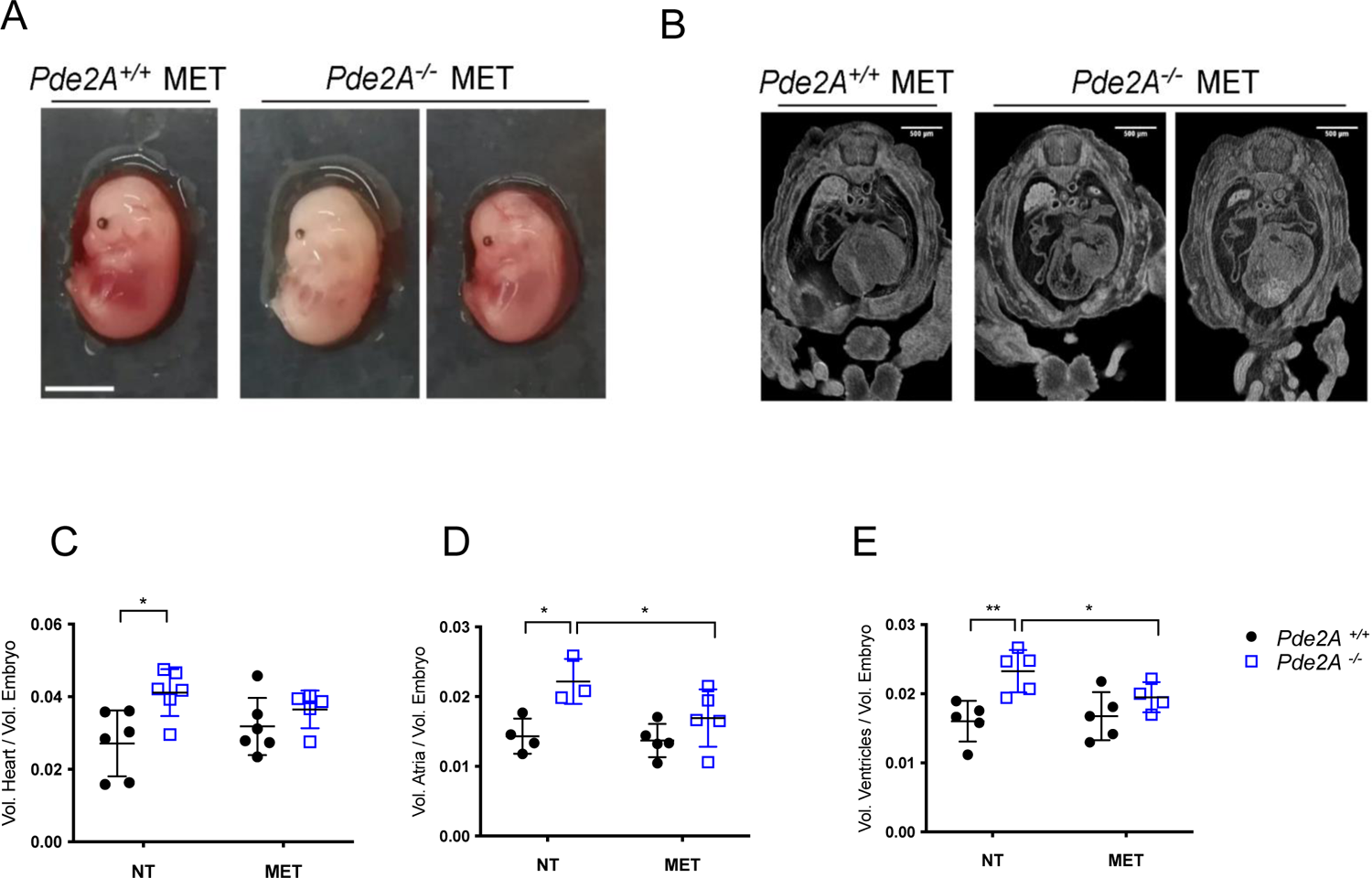
Micro-CT examination shows heart defect recovery after Metoprolol treatment. A) Picture of *Pde2A^+/+^* and *PDE2A^-/-^* embryos at E14.5 treated with Metoprolol. Scale bar=0.5cm. B) Micro-CT picture of the hearts in Metoprolol treated *Pde2A^+/+^* and *Pde2A^-/-^* embryos. C-D) Ratio between total (C), atrial (D), ventricular (E) volumes and embryo volumes obtained by micro-CT analyses of Metoprolol treated (MET) or not treated (NT) *Pde2A^+/+^* and *Pde2A^-/-^* embryos. At least n=4 for each genotype/treatment. ANOVA two-way was used to compare *Pde2A^-/-^* versus the relative *Pde2A^+/+^* and *Pde2A^-/-^* versus *Pde2A^-/-^* MET samples. * P≤0.05, ** P≤0.01, not significant values were not shown.

**Figure 5.**
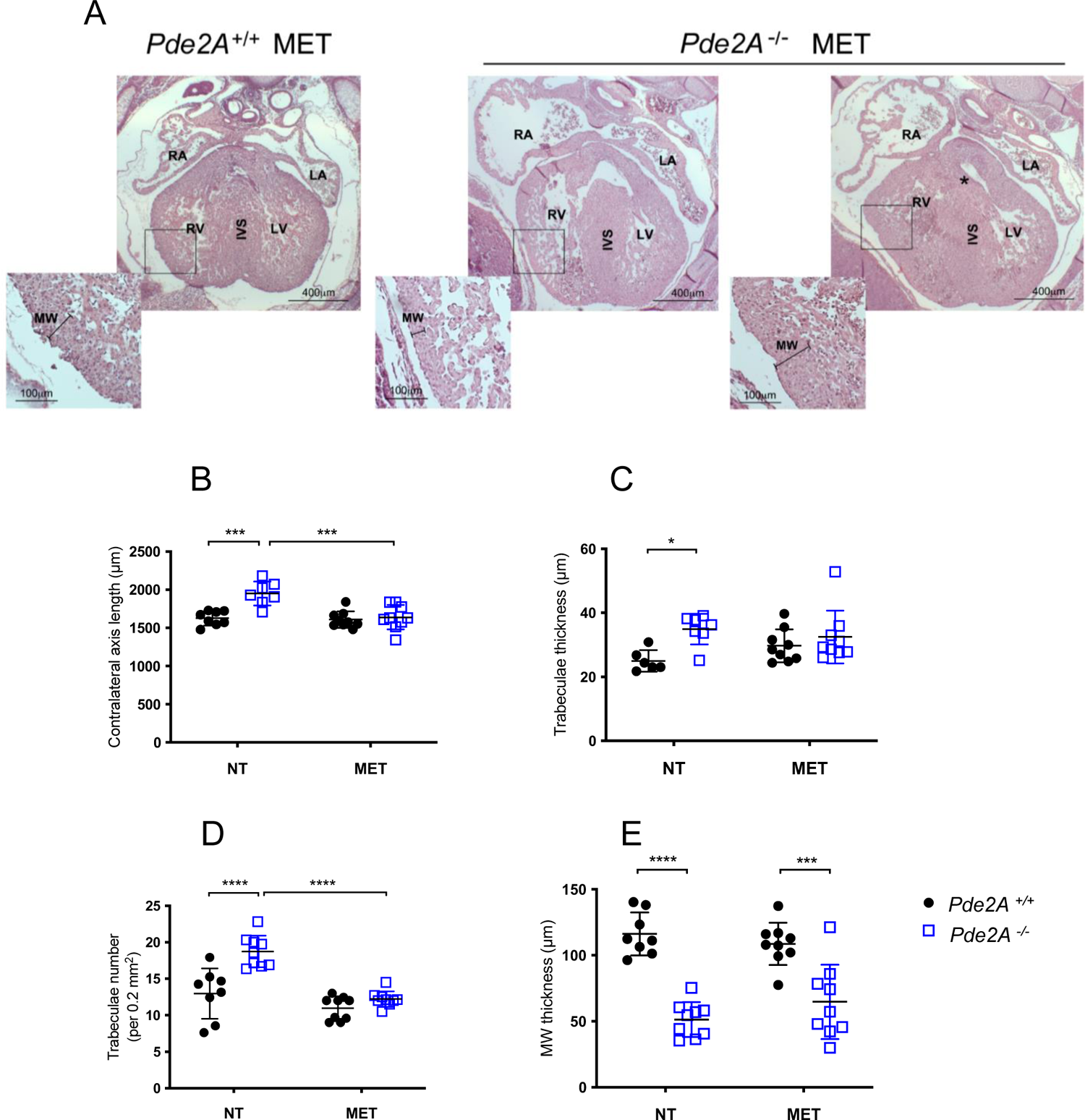
Histological analyses show heart defects recovery, except in myocardial wall, after Metoprolol treatment. A) Haematoxylin and Eosin staining of transversal sections of Metoprolol treated (MET) or not treated (NT) embryos, the heart is shown. Left and right ventricles (LV, RV), atria (LA, RA) and interventricular septum (IVS) are indicated. Insets show magnification of trabeculae and myocardial wall (MW). B-E) Graphs of contralateral axis (B), trabeculae thickness (C) and trabeculae number (D) and myocardial wall (E) evaluated by ZEISS AXIOSKOP 2 PLUS and ImageJ software. n=9 *Pde2A^+/+^* and n=11 *Pde2A^-/-^* NT embryos and n=4 *Pde2A^+/+^* and n=3 *Pde2A^-/-^* MET treated embryos; ANOVA two-way was used to compare *Pde2A^-/-^* versus the relative *Pde2A^+/+^* and *Pde2A^-/-^* versus *Pde2A^-/-^* NACMET samples. * P≤0.05, *** P≤0.001, **** P≤0.0001, not significant values were not shown.

### H89 treatment partially rescues CHD of *Pde2A^-/-^* embryos

To validate the cause-effect of the increase of cAMP level in the onset of CHDs, we used the PKA inhibitor H89. Morphological examination of H89 treated *Pde2A^-/-^* E14.5 embryos showed anemia, hemorrhages, and reduced liver size like not-treated embryos (Figure S4A). Micro-CT analysis revealed that H89 treatment was able to prevent the increase of total heart, atrial, and ventricular volumes observed in *Pde2A^-/-^* respect to *Pde2A^+/+^* embryos (Figures S4B-E). However, H89 *Pde2A^-/-^*treated hearts maintained ventricular septum defects (Figure S4B and Movie S4-S5).

As for Metoprolol treatment, heart microscopic examination of hematoxylin and eosin-stained embryonic sections revealed that H89 was able to prevent the increase of the contralateral axis, the increase of trabecular number and thickness, but not the ventricular septum defect or the reduction of the myocardial wall in *Pde2A^-/-^* (Figures S4F-L). These data straighten the hypothesis that inhibition of the cAMP/PKA system, although partially, is able to rescue CHDs of *Pde2A^-/-^* embryos.

### Metoprolol treatment restores the expression of oxidative stress genes in hearts of *Pde2A^-/-^* **embryos.**

We then evaluated the expression level of genes that resulted more upregulated in *Pde2A* knockout hearts by RNA-seq analysis and common to different pathways, in Metoprolol treated and untreated embryos. We observed that except for Fgf23, Metoprolol treatment did not affect the upregulation of Gata1 and Bmp10 found in *Pde2A^-/-^* as well as the expression increase of Epor, Ccl2, Ccl12 genes. (Figure 6A). Vice versa, the expression of several genes involved in oxidative stress, was reverted by Metoprolol treatment (Figure 6B). These results suggest that oxidative stress is counteracted by reestablishing the functional cAMP/PKA signaling.

**Figure 6.**
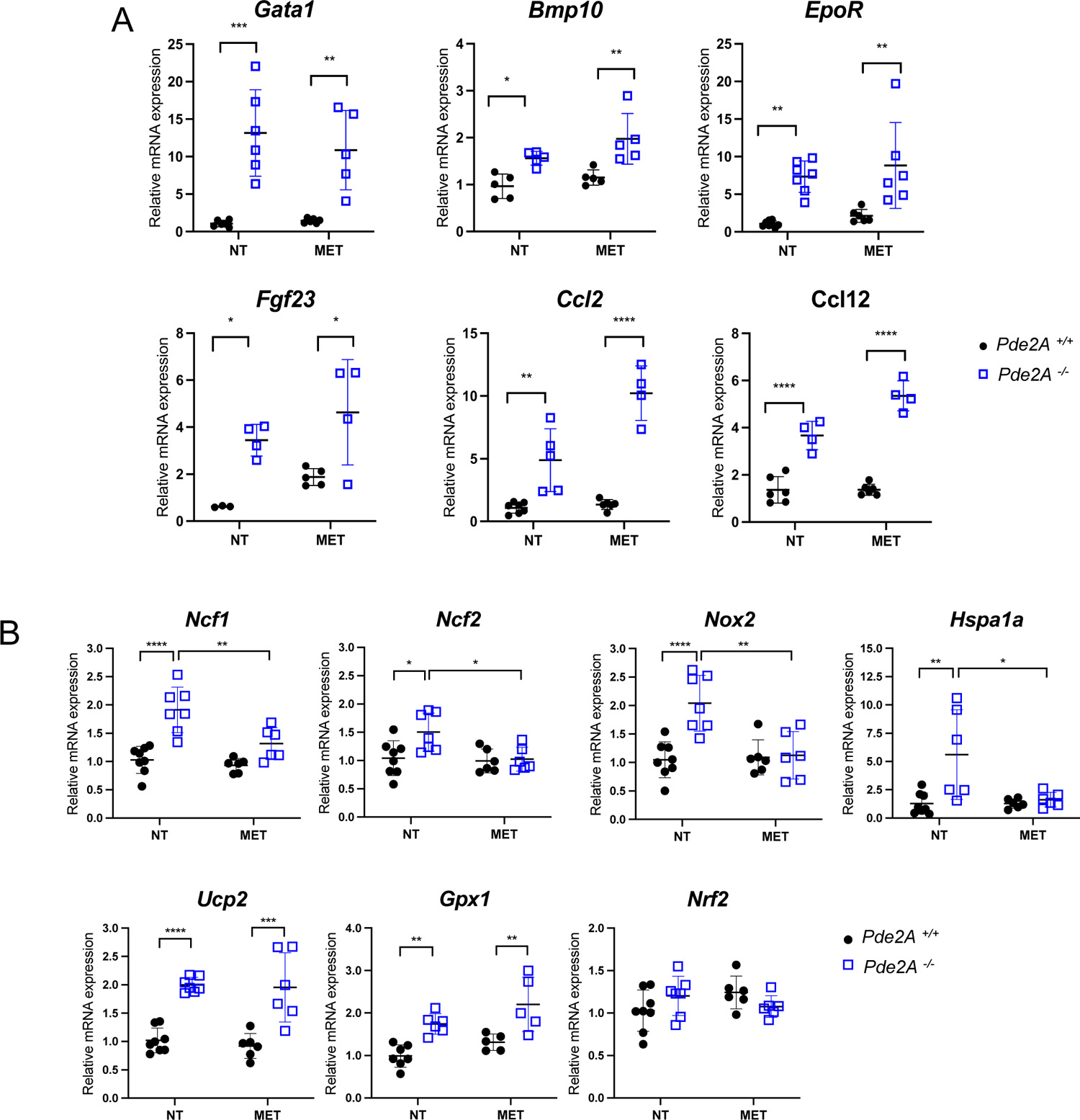
Prevention of oxidative stress gene up-regulation by Metoprolol treatment. A) Histograms of relative mRNA expression of most up-regulated genes involved in different biological processes evaluated by qRT-PCR. B) Histograms of relative mRNA expression of critical oxidative stress genes evaluated by qRT-PCR. n=9 *Pde2A^+/+^* and n=11 *Pde2A^-/-^* NT embryos and n=4 *Pde2A^+/+^* and n=3 *Pde2A^-/-^* MET treated embryos; ANOVA two-way was used to compare *Pde2A^-/-^* versus the relative *Pde2A^+/+^* and *Pde2A^-/-^* versus *Pde2A^-/-^* MET samples. * P≤0.05, ** P≤0.01, *** P≤0.001, **** P≤0.0001, not significant values were not shown.

### Treatment with the antioxidant NAC ameliorates cardiac defects in Pde2A^-/-^ embryos

Based on previous results indicating an involvement of the oxidative stress pathway in the heart defects of *Pde2A^-/-^* embryos, we checked whether these alterations could be rescued by NAC antioxidant treatment. Pregnant females were treated or not with NAC at the plug until the day before sacrifice. Phenotypically the mutant embryos appeared like the *Pde2A^-/-^* embryos (Figure 7A). The CM-H2DCFDA staining revealed that ROS positive cell increased in *Pde2A^-/-^* isolated cardiac cells, compared to *Pde2A^+/+^* cells confirming an increase of oxidative stress in the absence of Pde2A. NAC administration significantly counteracted ROS production in *Pde2A^-/-^* cardiac samples (Figure 7B).

**Figure 7.**
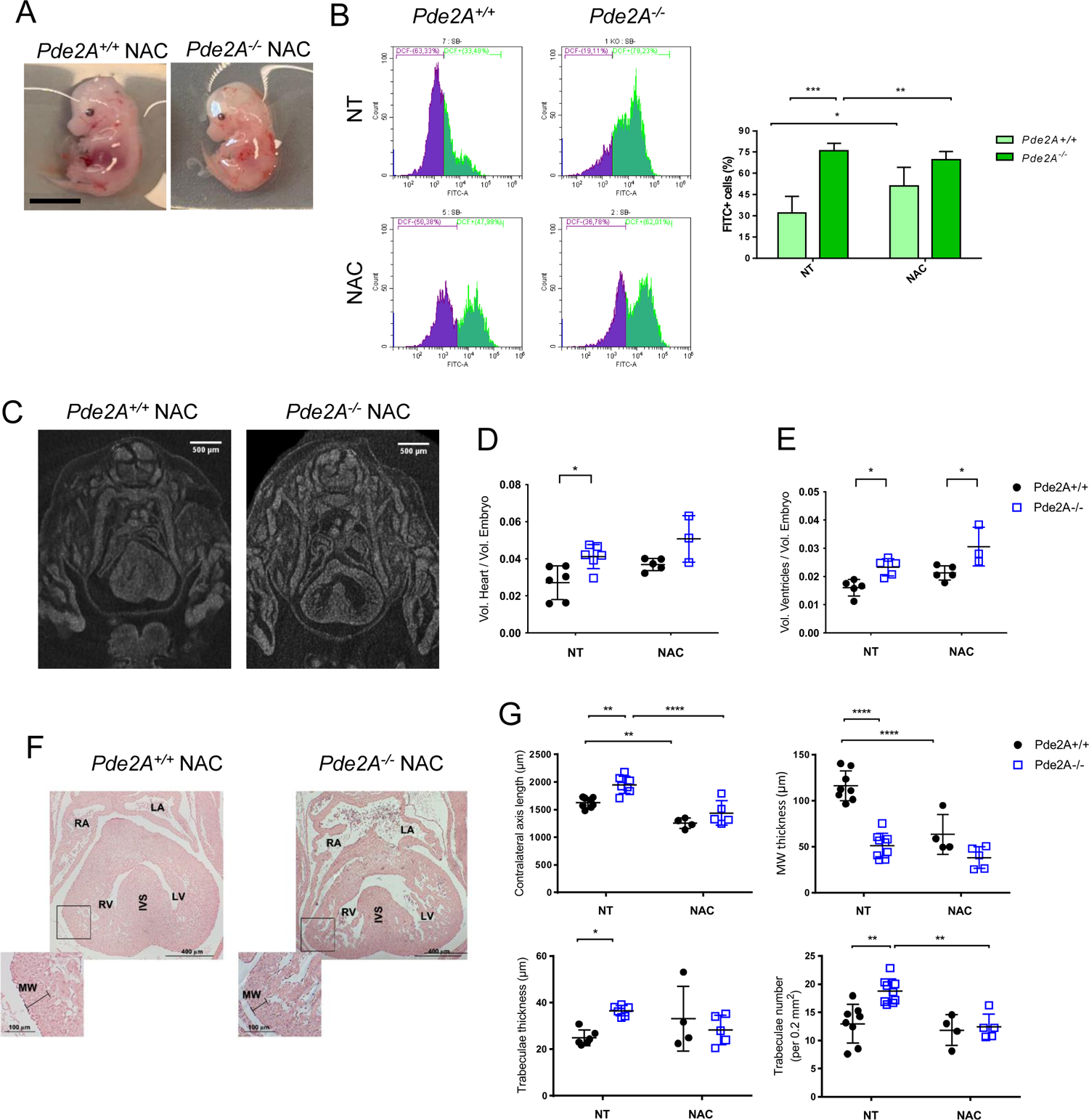
Oxidative stress and specific CHDs prevention by NAC administration. A) Picture of *Pde2A^+/+^* and *Pde2A^-/-^* embryos at E14.5 treated with NAC. Scale bar=0.5cm. B) Flow cytometry analysis of ROS level in cardiac cells isolated from *Pde2A^+/+^* and *Pde2A^-/-^* heart embryos in presence or absence of NAC. At least n=3 for each genotype and treatment. C) Micro-CT picture of the hearts in NAC treated *Pde2A^+/+^* and *Pde2A^-/-^* embryos. Scale bar=0.5cm. D-E) Ratio between total (D), ventricular (E) volumes and embryo volumes obtained by micro-CT analyses of NAC treated or NT *Pde2A^+/+^* and *PDE2A^-/-^* embryos. At least n=3 for each genotype /treatment. F) Haematoxylin and Eosin staining of transversal sections of NAC treated or NT embryos, the heart is shown. Scale bar=400μm. G) Graphs of contralateral axis, trabeculae thickness, trabeculae number and myocardial wall evaluated by ZEISS AXIOSKOP 2 PLUS and ImageJ software. At least n=6 *Pde2A^+/+^* and n=6 *Pde2A^-/-^* NT embryos and n=4 *Pde2A^+/+^* and n=5 *Pde2A^-/-^* NAC treated embryos; ANOVA two-way was used to compare *Pde2A^-/-^* versus the relative *Pde2A^+/+^*, *Pde2A^+/+^* versus *Pde2A^+/+^* NAC and *Pde2A^-/-^* versus *Pde2A^-/-^* NAC samples. * P≤0.05 ** P≤0.01, **** P≤0.0001, not significant values were not shown.

Micro-CT analysis did not reveal major rescue in *Pde2A^-/-^* embryos treated with NAC, except for total heart volume (Figures 7C-E and Movie S6-S7). However, histological analysis revealed improvement of parameters such as contralateral axis and trabecular number and thickness (Figure 7F-G). To be noted NAC appear to decrease some of these values also in *Pde2A^+/+^*embryos suggesting an effect on NAC in the basal homeostasis of cardiac oxidative stress during development.

Overall, these results indicate that antioxidants can be a valuable treatment for CHDs caused by cAMP unbalance.

## DISCUSSION

In this study RNA-seq was performed on isolated hearts from *Pde2A^+/+^*and *Pde2A^-/-^* at E14.5. Gene expression analysis revealed extensive alterations in the transcriptional profile of *Pde2A^-/-^* hearts, resulting in more than twenty thousand differentially expressed genes. Among these, 515 genes showed strong gene expression alteration. Notably, by performing a stringent analysis with DAVID functional enrichment, the transcriptome analysis revealed a deregulation of many pathways, including those involved in the immune response, angiogenesis, hypoxia, oxidative stress as well as transcription factors involved in the ERK signaling cascade. All these pathways are important for correct cardiac development and their alterations have also been reported in various cardiovascular diseases^33^.

Some of the genes found altered in *Pde2A^-/-^* hearts, such as Epor, Ccl2, and Fgf23 are described involved in mitochondrial metabolism other than many other functions^34–36^. The binding of erythropoietin to its receptor, Epor, activates endothelial nitric oxide synthase (eNOS) which is involved in the regulation of mitochondrial biogenesis, turnover and proliferation^34^. Ccl2 overexpression may be linked to alterations in mitochondrial dynamics that regulate anabolic and catabolic pathways, indicating a relationship between mitochondrial dysfunction, autophagy, and chronic disease^35^. High levels of Fgf23 are able to induce oxidative stress through the activation of NADPH oxidase complex^36^.

It is also well known that inflammatory cytokines and oxidative stress levels are observed in several cardiovascular diseases^37, 38^, however few inconclusive works have reported the role of inflammation and oxidative stress in CHDs in neonates and adults^39–42^.

Therefore, altogether the RNA-seq results indicated an alteration of mitochondrial function in the knockout model for *Pde2A*. A deeper investigation confirmed that genes associated with oxidative stress such as Nox2, Ncf1, Ncf2, Nrf2, Ucp2, Hspa1a, Gpx1 are significantly upregulated in *Pde2A^-/-^* embryonic hearts.

Nox2 is the catalytic subunit of the mitochondrial NADPH oxidase (Nox) that, upon activation, binds various regulatory subunits such as Ncf1 (neutrophil cytosolic factor 1) and Ncf2 (neutrophil cytosolic factor 2) and plays an important role in the production of ROS^43^. Physiologically, the cells have developed various mechanisms to protect themselves from the accumulation of ROS. These mechanisms include the prevention of the accumulation of ROS through uncoupling protein 2 (Ucp2), the reduction of hydrogen peroxide mediated by the nuclear factor erythroid 2-related factor 2 (Nrf2) and ROS degradation via glutathione peroxidase 1 (Gpx1)^44, 45^. In case of stress, it is also important to maintain proper protein homeostasis within the cell. Indeed, the chaperon heat shock protein 1A (Hspa1a), during cellular stress, might activate two opposite mechanisms, the folding, and the degradation of proteins depending on its acetylation/deacetylation state^46^.

The Metoprolol and H89 used in this study are drugs of clinical use that act upstream and downstream of the cAMP cascade, respectively. Both drugs were not able to prevent the gross phenotypes of *Pde2A* knockout embryos such as liver size, anemia, and hemorrhages.

However, it was found that embryos, whose mothers underwent Metoprolol treatment, recovered cardiac dimensions, hypertrabeculation defect and partially restore the interventricular septal defect. E14.5 embryos harvested from the mothers treated with H89, on the other hand, despite showing a recovery of cardiac dimension and hypertrabeculation, did not recover the interventricular septum. While significant differences continue to be present in the thickness of the ventricular wall, there is an improvement trend with both treatments. Molecular analyses carried out on hearts of E14.5 *Pde2A^-/-^*embryos showed that Metoprolol can reduce the level of cAMP and the expression of genes that induce oxidative stress (Nox2, Ncf1, Ncf2) whereas preserves the high expression of genes that protect from oxidative stress (Ucp2, Gpx1). Metoprolol treatment does not affect the upregulation of genes implicated in inflammatory response possibly because themselves trigger oxidative stress or because the left heart defects maintain the inflammatory state.

NAC is a safe and well tolerated compound displaying antioxidant properties; it acts mainly by providing the intracellular cysteine necessary to increase glutathione levels. Our results indicate that NAC attenuated just some CHDs carried by *Pde2A^-/-^* embryos, possibly because of the low drug bioavailability from the mother to the embryos. In addition, NAC, despite having reduced ROS in *Pde2A^-/-^* cardiac cells, slightly increased the global oxidative stress in the *Pde2A^+/+^* hearts. Dose increase, modification of drug administration procedures, or combination of NAC with other antioxidants and/or anti-inflammatory compounds may be more effective in preventing CHDs.

This study highlighted the involvement of Pde2A in new biological processes such as inflammation, angiogenesis, transcription and oxidative stress and how pharmacological treatments aimed at modulating cAMP levels can counteract congenital heart defects, in part by restoring the oxidative stress response. It is not yet clear to what extent the different types of cells are differentially affected in the heart of *Pde2A^-/-^* versus *Pde2A^+/+^*embryos and how the used drugs impact on these cell populations. Future studies will address this topic by using flow cytometry and spatial digital transcriptomic analyses. To date, non-human CHDs related to the complete absence of PDE2A has been found, probably because, as in the mouse model, homozygotes are embryonic lethal. However, one report, describing a patient with a homozygous inactivating mutation in the PDE2A^47^ gene, has been published. The patient suffers from a form of Huntington’s chorea, but it will be worth to evaluate its cardiac performance with electrocardiograms, arrhythmias, and cardiac function.

Overall, the information obtained from our study will be useful to investigate human CHDs associated with the compromission/imbalance of the cAMP/PDE2A system and to investigate therapies suitable for their recovery.

## Nonstandard abbreviation and acronyms

PDE: phosphodiesterase

CHD: Congenital Heart Disease

NAC: N-Acetyl-Cysteine

ROS: Reactive Oxygen Species

Micro−CT: micro-Computed Tomography

PKA: Protein kinase A

DCM: Dilated Cardio Myopathy

CM-H2DCFDA: 2’,7’-dichlorodihydrofluorescein diacetate

RNA-seq: RNA sequencing

NADPH: Nicotinamide Adenine Dinucleotide Phosphate

## Acknowledgements

We thank EMMA service project that founded the EC FP7 Capacities Specific Program and allow to obtain *Pde2A^+/-^*mice. We are particularly thankful to Dr. Vladimir Benes, EMBL Head of Genomics Core Facility, EMBL Heidelberg, Germany for the RNA-seq processing, Dr. Emerald Perlas, Head of Histology Core Facility, EMBL Rome, Italy for histology suggestions, to Prof. Mauro Giorgi, University of Sapienza, Italy and Dr. Giovina Ruberti, IBBC-CNR, Monterotondo Rome, Italy for critical reading.

## List of Supplemental materials

Table S1

Table S2

Figure S1-S4

Video Movie S1-S7

Major Resources Table

